# Spatio-temporal regulation of ligand trafficking and TLR9 activation involves PIEZO1 mechanosensing in human plasmacytoid dendritic cells

**DOI:** 10.1101/2025.07.31.667789

**Authors:** Shrestha Pattanayak, Deblina Raychaudhuri, Purbita Bandopadhyay, Upasana Mukhopadhyay, Tithi Mandal, Chinky Shiu Chen Liu, Nidhi Kalidas, Sumangal Roychowdhury, Krishnananda Chattopadhyay, Fnu Ashish, Bidisha Sinha, Dipyaman Ganguly

## Abstract

Plasmacytoid dendritic cells (pDCs) are specialized innate immune cells which play a pivotal role in antiviral immunity by producing large quantities of type I interferons (IFNs) upon sensing nucleic acids via Toll-like receptor 9 (TLR9). Synthetic oligodeoxynucleotides (ODNs) such as CpGA and CpGB, both containing unmethylated CpG motifs, are commonly used experimental TLR9 agonists. Interestingly, CpGA and CpGB elicit markedly different responses in pDCs – CpGA induces robust IFN-α production, whereas CpGB does not. The mechanistic basis underlying this ligand-specific functional divergence has remained unclear. Here, we identify PIEZO1, a mechanosensory ion channel, as a critical determinant of ligand-specific IFN responses in human primary pDCs. We demonstrate that unlike CpGB, CpGA self-associates into larger aggregates that generate membrane tension during cellular uptake, leading to activation of PIEZO1. This activation triggers localized calcium influx and cytoskeletal remodeling, resulting in the formation of local F-actin structures that retain CpGA within early endosomes enabling sustained IRF7 nuclear translocation and robust type I IFN production. Disruption of PIEZO1 or actin polymerization abrogates CpGA-induced IFN production, while pharmacological activation of Piezo1 enhances IFN production in response to CpGB. Thus, these findings uncover a previously unrecognized biophysical checkpoint in nucleic acid sensing, wherein membrane tension is transduced via PIEZO1 into spatially controlled TLR9 signaling. Overall, our study establishes PIEZO1 mechanosensing at the plasma membrane as a key regulatory event in nucleic acid-induced immunity, with different functional outcomes based on the cargo structure, opening potential new avenues for modulating type I IFN responses in infection and autoimmunity.

## Introduction

Plasmacytoid dendritic cells (pDCs) are a subset of innate immune cells specialized for the rapid and abundant production of type I interferons (IFNs) in response to nucleic acids (Liu, 2005; Reizis, 2019). This capacity is predominantly mediated through the endosomal recognition of unmethylated CpG motifs in DNA by Toll-like receptor 9 (TLR9) (Gilliet et al, 2008; Swiecki et al, 2015). Synthetic oligodeoxynucleotides (ODNs) bearing CpG motifs have been widely employed to dissect pDC responses and are broadly classified into three categories-CpGA, CpGB, and CpGC, based on their structural modifications and immunostimulatory profiles (McKenna et al, 2005). Among these, CpGA is particularly potent at inducing type I IFNs, whereas CpGB predominantly triggers proinflammatory cytokine (TNF-α and IL-6) production, but elicits a markedly blunted type I IFN response (Kerkmann et al, 2003; McKenna et al, 2005; Chinnathambi et al, 2012).

Thus, despite comparable binding affinities for TLR9, CpGA and CpGB elicit fundamentally distinct signaling outcomes, raising a longstanding and unresolved question: how do chemically related ligands engaging the same receptor drive divergent transcriptional programs in pDCs? Existing literature suggests that endosomal trafficking plays a central role in this dichotomy. CpGA is retained in early endosomes, where they interact with TLR9 driving IRF7 activation and type I IFN production, while CpGB is trafficked more rapidly to late endolysosomes, where it predominantly activates NF-κB (Honda et al, 2005; Guiducci et al, 2006). This differential response based on ligand characteristic is interesting, more so because of inherent property of CpGA to form multimeric aggregates as opposed to CpGB (Guiducci et al, 2006; Chinnathambi et al, 2012). Similar endo-lysosomal sorting has also been documented with immune complexes generated in the context of autoimmune diseases (Lande et al, 2007; Ganguly et al, 2009; Ganguly et al, 2018). However, the upstream mechanisms that regulate ligand sorting and differential spatial compartmentalization of TLR9 signaling to induce type I IFN production remain poorly defined.

To address this, we initially investigated whether CpGA and CpGB differ in their modes of cellular entry or early endocytic uptake pathways in pDCs. Surprisingly, we did not find any significant differences in uptake efficiency or route of endocytosis between the two ligands. However, scanning electron microscopy revealed a striking difference: CpGA stimulation, unlike CpGB, resulted in pronounced membrane ruffling. Plasma membrane ruffling following internalization of bacteria has been demonstrated to induce membrane tension in local plasma membrane, which is sensed by PIEZO1 mechanosensors (Tadala et al, 2022). Thus, the membrane ruffling observed in response to CpGA uptake in human pDCs made us hypothesize a similar possibility.

We considered whether differences in the biophysical properties of the ligands, such as aggregation and cargo size, might be sensed by pDCs during uptake, and contribute to their distinct functional outcomes. Specifically, we hypothesized that CpGA and CpGB may differentially engage cellular mechanotransduction pathways during internalization, thereby directing distinct intracellular trafficking and signaling outcomes. The mechanosensitive ion channel PIEZO1 has emerged as key sensors of membrane deformation and tension, regulating diverse biological processes ranging from vascular development to epithelial cell integrity (Murthy et al, 2017). Our previous work delineated the critical role of PIEZO1 in regulating human T cell activation, function and migration (Liu et al, 2018; Liu et al, 2019; Liu et al, 2024). However, whether mechanotransduction via PIEZO1 plays any role in innate immune sensing of nucleic acids and type I IFN response remains entirely unexplored.

Here, we identify PIEZO1 as a key mechanosensory gatekeeper that enables human pDCs to discriminate between CpG ligands based on their physical properties. We demonstrate that CpGA, but not CpGB, forms large self-associated molecular aggregates that result in greater increase in plasma membrane tension during cellular entry. This tension is sensed by PIEZO1, which becomes rapidly activated and clustered at sites of CpGA uptake, leading to localized calcium influx. In turn, PIEZO1-mediated calcium signaling drives F-actin polymerization around early endosomes, forming a stabilizing meshwork that retains CpGA in these signaling-permissive compartments and sustains IRF7 activation and nuclear translocation. Genetic or pharmacological inhibition of PIEZO1 disrupts this axis, resulting in premature early endosomal escape of CpGA, impaired IRF7 nuclear translocation, and a sharp reduction in IFN-α production. Strikingly, chemical activation of PIEZO1 is sufficient to enhance IFN production in response to CpGB, highlighting a ligand-independent mechanism to tune Type I IFN responses in pDCs.

Collectively, our findings reveal that PIEZO1-mediated mechanosensation serves as a critical upstream regulator of CpGA trafficking and type I IFN production in pDCs. This study provides the first evidence that innate immune recognition of nucleic acids is not solely determined by ligand chemistry or receptor specificity, but also by the mechanical forces generated during antigen encounter. By coupling membrane tension to peri-endosomal actin remodeling and transcriptional output, PIEZO1 enables pDCs to convert physical properties of nucleic acid ligands into spatially and temporally precise cytokine responses. Thus, these insights not only address the long-standing question of CpG class-specific signaling but also perhaps highlight a more general principle by which innate immune cells interpret the physical properties of ligands to fine-tune cytokine output. Overall, the findings from this study not only resolve a key mechanistic gap in our understanding of TLR9 activation in human pDCs, but also indicate possible new directions to modulate pDC activation and type I IFN-driven antiviral immunity and autoimmunity through the biophysical tuning of the structural attributes of TLR ligands or combination therapies targeting the PIEZO1 mechanosensors.

## Results

### Self-association prone CpGA ligands generate greater membrane tension than CpGB during internalization by primary human pDCs

To investigate how the physical and structural properties of synthetic TLR9 ligands influence their uptake and downstream signaling in pDCs, we first compared CpGA and CpGB in terms of their supramolecular organization in solution. Analyzing Small-angle X-ray scattering (SAXS) and fluorescence correlation spectroscopy (FCS) data, we found that CpGA spontaneously forms high-molecular-weight aggregates in PBS, whereas CpGB remains predominantly monomeric or assembles into only small oligomers (Figure 1A, B), consistent with previous reports (Guiducci et al, 2006; Chinnathambi et al, 2012). In their SAXS profiles, the downward trend of data points as s values approached to 0 nm^-1^ confirmed that both CpGA and CpGB molecules adopt pronounced interparticulate effect in water (Supplementary Figure 1A and B). This phenomenon is known for charged molecules as they repel each other in solution causing them to appear smaller than their usual hydrodynamic profile (Stradner et al, 2004; Jacques et al, 2010; Jacques et al, 2012). This effect occurs mainly in the absence of counter ions, and it disappeared completely when the samples were moved to PBS. Interestingly, the increment in temperature did not change the solution scattering profile of the molecules in either solvent indicating that the solution shape of both CpGA and CpGB is stable in the solvents, with or without counter ions. Presuming globular shape profile, the Guinier approximation of the averaged datasets provided R_g_ values of 5.63 and 2.65 nm for CpGA and CpGB in PBS, respectively. Comparison with their values in water suggested that in PBS, CpGA molecules self-associate significantly while CpGB molecules opens up slightly. Guinier analysis for rod-like shape provided R_c_ values of 0.78 and 0.80 nm for CpGA and CpGB in PBS, respectively. Deduced parameters provided L and A values of 19.31 and 8.75 nm, and 7.22 and 3.31 for CpGA and CpGB in PBS, respectively. This clearly indicated differential behavior of solution shape and self-association order for CpGA molecules, which appeared to now form exceptionally elongated shape with very skewed aspect ratio compared to CpGB molecules. Dimensionless Kratky plots for both molecules peaked close to sR_g_ value of 1.73 (dotted lines in middle panels of Supplementary Fig 1A, B) which suggested disordered globular shape (Rambo et al, 2010; Rambo et al, 2011). Bayesian inference of the SAXS datasets indicated average molecular weight of 53.1 and 9.5 kDa for CpGA and CpGB molecules in PBS. Molecular envelope shapes were calculated for both CpGA and CpGB molecules in PBS and water (Figure 1A and Supplementary Figure 1C). Notably, the minimal aggregate size of CpGA was more than twice that of CpGB in PBS, demonstrating substantial differences in particle size and structural complexity between the two ligands (Figure 1A). However, there was no difference in the size of CpGA and CpGB in water (Supplementary Figure 1C).

**Figure 1:**
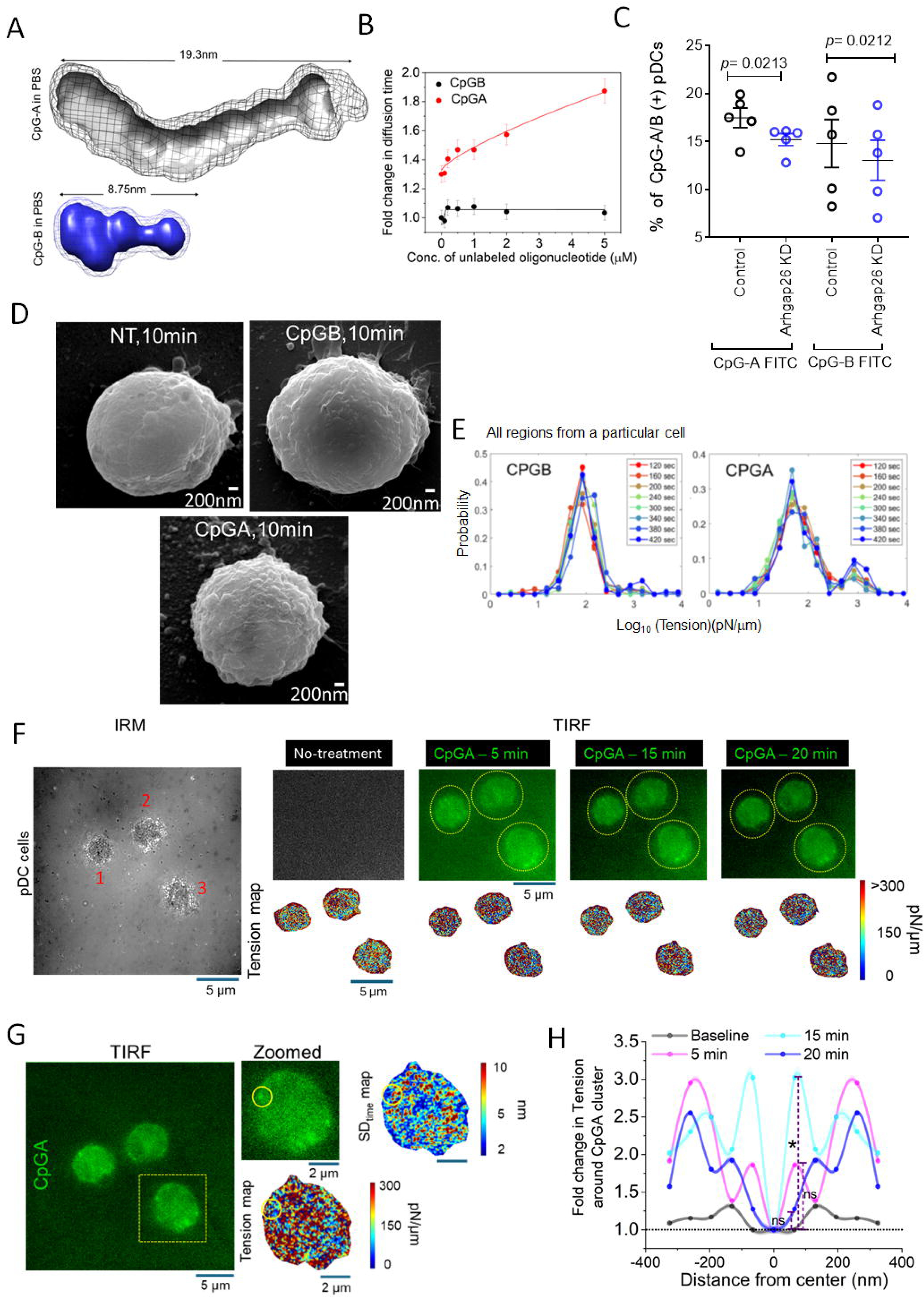
Self-associated CpGA aggregates generate greater membrane tension than CpGB during internalization by primary human pDCs. **A)** Molecular maps of the envelope shapes, calculated for CpGA (grey) and CpGB (blue) molecules in PBS, are presented. Solid surface represents the common shape in ten restorations and mesh around represents the variation in the individual models computed. Numbers mentioned are measured approximate end-to-end dimensions in nm, and correlate with persistence length calculated from respective SAXS datasets. **B)** Plot of fold change in the diffusion time (with respect to 20nM Alexa-488 labeled CpGB) vs. concentration of unlabeled oligonucleotide (CpGA in red and CpGB in black) observed in FCS experiment. CpGB did not show any significant change in the diffusion time over the indicated range whereas CpGA showed a change in diffusion time indicates self association. Three independent measurements were taken (n=3). The data were plotted as mean ± SD **C)** Flow cytometric analysis of Alexa fluor-488 labeled CpGA or CpGB uptake in control and ARHGAP26 knockdown pDCs (n=5). Data represent mean ± s.e.m from 3 independent experiments. Statistical significance was assessed using Student’s t-test. **D)** Representative scanning electron microscopy images demonstrating membrane morphology in untreated pDCs, or cells stimulated with CpGA or CpGB for 10 minutes. Images are representative of 4 independent experiments. **E)** Evolution of tension distribution for different regions of a particular cell for two minutes followed in time after treatment with CpGB/A. **F)** Representative IRM image of cells followed in time before and after CpGA-FITC addition with their corresponding TIRF fluorescence images attached. The pixel wise tension maps of the cells along time are also represented. Scale bar-5 µm. **G)** A typical TIRF frame with cells treated with CpGA-FITC. Scale bar-5 µm. The dotted yellow rectangular line marks the cell that is zoomed in. The zoomed in view shows a CpGA cluster with its corresponding pixel-wise tension and SDtime maps. Scale bar-2 µm. The yellow circle marks the cluster. **H)** Radial profiles of Tension around the Fluorescence peaks of CpGA clusters. Error denoted by the shaded region as SEM. * - represents significant change from the baseline condition for that particular time point after CpGA addition within 100 nm from center. Mann–Whitney U statistical significance test where ns-p >0.05, * - p <0.05, ** - p <0.001. The dotted black line through the plots denotes the value ‘1’. For CpGA, Ncell: 9; Ncluster: 46.

Next, to assess the self-association propensity between CpGA and CpGB oligonucleotide we used single molecule fluorescence-based Fluorescence Correlation Spectroscopy (FCS). FCS measures the fluctuation of fluorescence intensity in a small detection volume (∼fl) to determine the dynamics of the biomolecules (Ray et al, 2025). We used a concentration dependent study by using 20nM Alexa-488 labeled oligonucleotide (CpGA or CpGB) with increasing concentration of unlabeled oligonucleotide (CpGA or CpGB) upto 5µM. In the case of CpGB, we did not observe any significant shift in the normalised autocorrelation function (Supplementary Figure 1D) and in the fold change in the diffusion time obtained from FCS measurements (Figure 1B). Any change in the number of particle (N) present in the confocal volume indicates association or dissociation of the detected species. Decease in the number of particles inside the detection volume indicates self-association or oligomerisation of the species under consideration (Ghosh et al, 2018; Das et al, 2023). The number of particle (N) analysis from FCS curve fitting using 3D-gaussian single diffusing species did not show any significant change for the entire concentration change (Supplementary Figure 1F). All together, these confirmed the absence of any higher order species formed by CpGB in the solution. On the contrary, for CpGA, we observed a shift in the normalised autocorrelation function (Supplementary figure 1E) and also a significant change of the fold change in the diffusion time (Figure 1B). With 1μM unlabeled CpGA concentration, the number of monomers associated is ∼3 (assuming a cube root dependence of diffusion with molecular weight) and with 5μM unlabeled CpGA concentration, the number of monomers associated is ∼6. The number of particle (N) analysis for CpGA showed a significant decrease indicating self-association (Supplementary Figure 1F).

Engagement of different endocytic pathways (viz. clathrin-mediated vs caveolin-mediated) have been shown to be regulated by the size of the cargo as shown earlier using different sized microspheres (Rejman et al, 2004). This raised a critical question: do these differences in aggregation state and particle size contribute to their divergent functional outcomes in pDCs by engaging distinct endocytic pathways? To address this, we used pharmacological inhibitors targeting clathrin- and caveolin-mediated endocytosis-Pitstop 2 and Genistein, respectively (Smith et al, 2013; Zhang et al, 2018). Inhibition of these classical uptake routes did not significantly impair internalization of either CpGA or CpGB (Supplementary Figure 2A), indicating that both ligands are likely internalized through non-canonical pathways. We next focused on the clathrin and caveolin-independent CLIC-GEEC pathway and found that siRNA-mediated knockdown of ARHGAP26 (Supplementary Figure 2B), the driver adaptor molecule of CLIC-GEEC endocytosis (Lundmark et al, 2008), led to reduced uptake of both CpGA and CpGB (Figure 1C, Supplementary Figure 2C). These findings indicated that the CLIC-GEEC pathway is involved in the internalization of both ligands and that differences in their functional outcomes are unlikely to be driven by differential endocytic routing. Interestingly, scanning electron microscopy of primary human pDCs treated with CpGA or CpGB for 10 minutes demonstrated striking differences in membrane morphology. CpGA-treated cells exhibited markedly more membrane ruffling compared to CpGB-treated cells (Figure 1D). This observation led us to hypothesize that the internalization of the larger CpGA cargo leads to greater membrane deformation, thereby imposing greater membrane stretch and mechanical strain on the plasma membrane, potentially serving as a mechanical cue.

### Increased local membrane tension surrounding endocytic pits during uptake of CpGA by human pDCs

To determine whether the larger size of self-associated aggregates of CpGA induces a global increase in the basal cell membrane tension during uptake, we performed Interference Reflection Microscopy (IRM) on cells before and after administration of CpGs. IRM is a non-invasive live cell imaging technique to quantify the nanometric height fluctuations of the basal membrane of adherent cells (Biswas et al, 2017). The characteristics of the height fluctuations of the membrane are primarily determined by its mechanical state. Hence, by analyzing spatio-temporal patterns of the height fluctuations, especially by fitting the power spectral density (PSD) with Helfrich-based model, estimates of the mechanical parameters, including fluctuation tension (Shiba et al, 2016) – the effective tension of the basal membrane, could be derived as previously reported (Biswas et al, 2017). On following same cells after addition of CpGA/B (Supplementary Video 1), we observed that CpGA but not CpGB increased the probability of finding higher tensed membrane patches in the cells. This is symbolized by the second peak of probability at higher tension values visible for CpGA but not for CpGB (Supplementary Figure 2D). The amplitude of the height fluctuations was quantified by measuring the standard deviation of relative height from time-series of height fluctuations (termed SD_time_). There was a concomitant pronounced decrease in SD_time_ as tension increased (Supplementary Figure 2E). Further, following single cells through time clearly revealed the building of the second peak in tension with time (Figure 1E) indicating that certain regions within CpGA-treated cells had enhanced tension rather than only the average tension being increased.

Therefore, we next focussed on understanding the impact on the membrane close to the cluster of CpGA. We utilized a sequential correlative imaging approach combining Total Internal Reflection Fluorescence (TIRF) microscopy and Interference Reflection Microscopy (IRM) (Supplementary Figure 3A) and imaged cells over time (Supplementary Figure 3B). TIRF enabled imaging of fluorescently labeled CpGA close to the basal membrane – due to low penetration depth of the evanescent-mode, while IRM enabled mapping of the effective tension (Figure 1F) and SD_time_ (Supplementary Figure 3C) in these single cells over time. Particular clusters were identified (Figure 1G) through TIRF images and maps of tension and SD_time_ generated from IRM images. To quantify the local effect, line scans of the tension, SD_time_ and fluorescence profiles were performed along x and y directions, centered at fluorescence peaks (indicating center of the cluster) (Supplementary Figure 3D). The averaged normalized profiles of tension around the fluorescence peak depict the fold change in tension as one moved away from the center. Such profiles were plotted for the 5-, 15-, and 20-minute time points (Figure 1H) along with profiles of the fluorescence at those regions (Supplementary Figure 3E). The baseline (or untreated) (Figure 1H) represents the state of tension at the same region where CpGA later clusters after its addition. By 15 minutes, we observed a distinct increase in membrane tension within approximately 100 nm of the CpGA cargo compared to the central region (Figure 1H). Although membrane tension here is dependent on the form of Helfrich model chosen to fit the fluctuation spectra, we demonstrate that direct measurements of membrane fluctuation amplitudes also fully reflect the impact of local membrane remodeling by CpGA, as the increase in effective tension was accompanied by a decrease in SD_time_ (Supplementary Figure 3E). This supports the idea that CpGA uptake locally elevates membrane tension around its site. From the cluster center, the maximum surge in fluctuation tension produced by CpGA cargo, within 325 nm, was compared with the untreated condition. We find the maximum tension increase (difference of tension at pixels from tension at center of CpGA cluster) to be lower when untreated while peaking at ∼ 1300 pN/µm at 5 and 15 min after CPGA addition (Supplementary Figure 3F). Interestingly, Piezo activation has been known to be triggered at a tension range of ∼ 600 - 2100 pN/µm (Lewis et al, 2015; Cox et al, 2016; Nourse et al, 2017; Wu et al, 2017; Yang et al, 2022; Lewis et al, 2024), although the tension threshold is expected to be system-dependent. The tension increase measured in this data (Supplementary Figure 3F) at 15 min in comparison to baseline (Supplementary Figure 3G), thus, could be sufficient to trigger PIEZO1 activation.

Overall, these findings suggested that the differential mechanical impact of CpGA versus CpGB is a function of the cargo size and resulting membrane tension, rather than receptor-specific endocytic sorting pathways.

### CpGA-induced membrane tension activates PIEZO1 to selectively enhance IFN-a production by human pDCs

Given the pronounced localized increase in membrane tension during CpGA exposure, we hypothesized that the mechanosensitive ion channel PIEZO1 may serve as a mechanosensor linking ligand size to immune signaling. Consistent with our hypothesis, confocal microscopy revealed rapid membrane recruitment and clustering of PIEZO1 in pDCs following CpGA, but not CpGB treatment (Figure 2A) suggesting that only CpGA induces sufficient membrane tension to cause PIEZO1 clustering. Additionally, CpGA triggered significantly higher levels of IFN-α compared to CpGB in primary human pDCs (Figure 2B) consistent with previous reports. Therefore, to assess whether PIEZO1 activation contributes to this differential type I IFN response in pDCs, we quantified IFN-α secretion following PIEZO1 inhibition. Pharmacological inhibition with GsMTx4 (Bae et al, 2011, Gnanasambandam et al, 2017) or siRNA-mediated knockdown of *PIEZO1* (Supplementary Figure 3H) significantly reduced CpGA-induced IFN-α levels at both the mRNA and protein levels (Figure 2C-E). Further, to test whether PIEZO1 activation is essential to promote a CpGA-like interferon response, we co-stimulated CpGB-treated cells with Yoda1, a selective PIEZO1 agonist (Syeda et al, 2015; Botello-Smith et al, 2019; Goon et al, 2024). Yoda1 markedly enhanced CpGB-induced IFN-α production (Figure 2F), and this increase was abrogated in PIEZO1 knockdown pDCs (Figure 2G, H). These results demonstrated that PIEZO1 activation plays a critical role in driving type I IFN responses in pDCs. Together, these findings implicated PIEZO1 as a mechanosensor that directly links biophysical properties of TLR9 ligands to type I IFN production in human pDCs.

**Figure 2:**
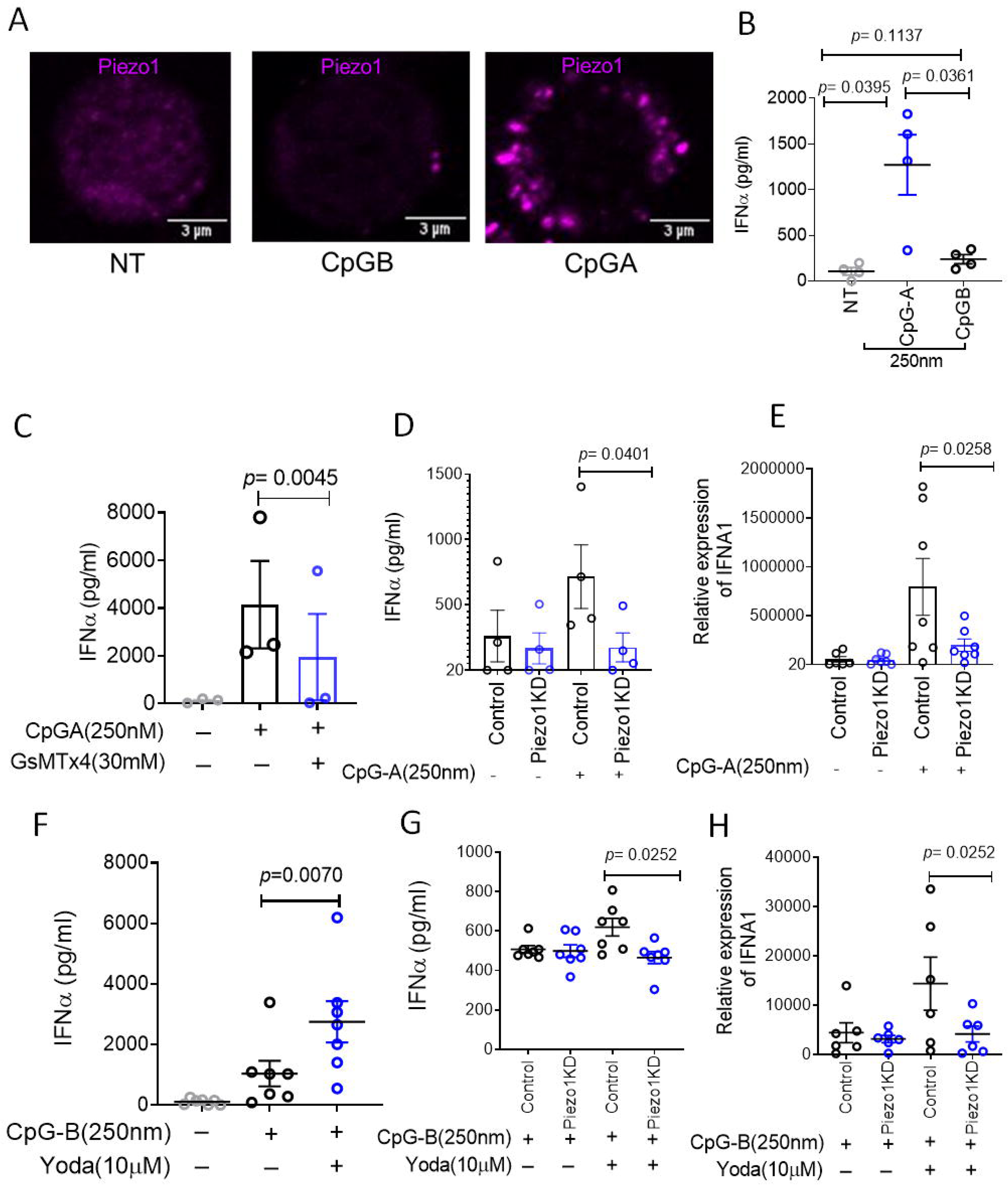
CpGA-induced membrane tension activates PIEZO1 to selectively enhance IFN-a production by pDCs. **A)** Representative confocal microscopy images showing intracellular PIEZO1 distrubution in untreated pDCs and those treated with CpGA or CpGB. **B,C)** IFNα production in pDC culture supernatants following stimulation with CpGA or CpGB **(B)**, and CpGA with increasing doses of GsMTx4 **(C)**, measured by ELISA. n=3-4. **D, E)** IFNα secretion **(D)** and *IFNA1* mRNA expression **(E)** in control and PIEZO1-knockdown (PIEZO1 KD) pDCs treated with or without CpGA. n=4-7. **F)** IFNα production by pDCs stimulated with CpGB in the presence or absence of Yoda1. n=7. **G, H)** IFNα secretion **(G)** and *IFNA1* gene expression **(H)** in control and PIEZO1 KD pDCs stimulated with CpGB with or without Yoda1 was measured. n=6-7. Data represent mean ± s.e.m. Statistical analysis was performed using Student’s t-test.

### PIEZO1 activation promotes CpGA retention in early endosomes and IRF7 nuclear translocation in human pDCs

We next investigated how PIEZO1 activation enhances type I IFN production by modulating the subcellular trafficking of CpGA. To determine the intracellular localization of CpGA, we performed confocal microscopy using markers for early endosomes (EEA1) and late endosomes (LAMP1). Consistent with previous studies, CpGA predominantly localized to early endosomes in primary human pDCs (Figure 3A, B, Supplementary Videos 2-3). Notably, this early endosomal retention of CpGA was dependent on PIEZO1 activity. Pharmacological inhibition of PIEZO1 using GsMTx4 significantly decreased CpGA colocalization with EEA1 and increased its colocalization with LAMP1 (Figure 3A, B, Supplementary Videos 4-5), suggesting a faster transition of CpGA to late endosomes and reduced residence time in early endosomes. Further, to confirm the critical role of PIEZO1 in regulating TLR9 ligand trafficking in pDCs, we co-treated CpGB-stimulated pDCs with Yoda1. Subcellular fractionation analysis showed that Yoda1 enhances CpGB accumulation in early endosomes (Figure 3C), supporting the role of PIEZO1 in promoting retention of TLR9 ligands in early endosomal compartments, thereby prolonging their signaling potential.

**Figure 3:**
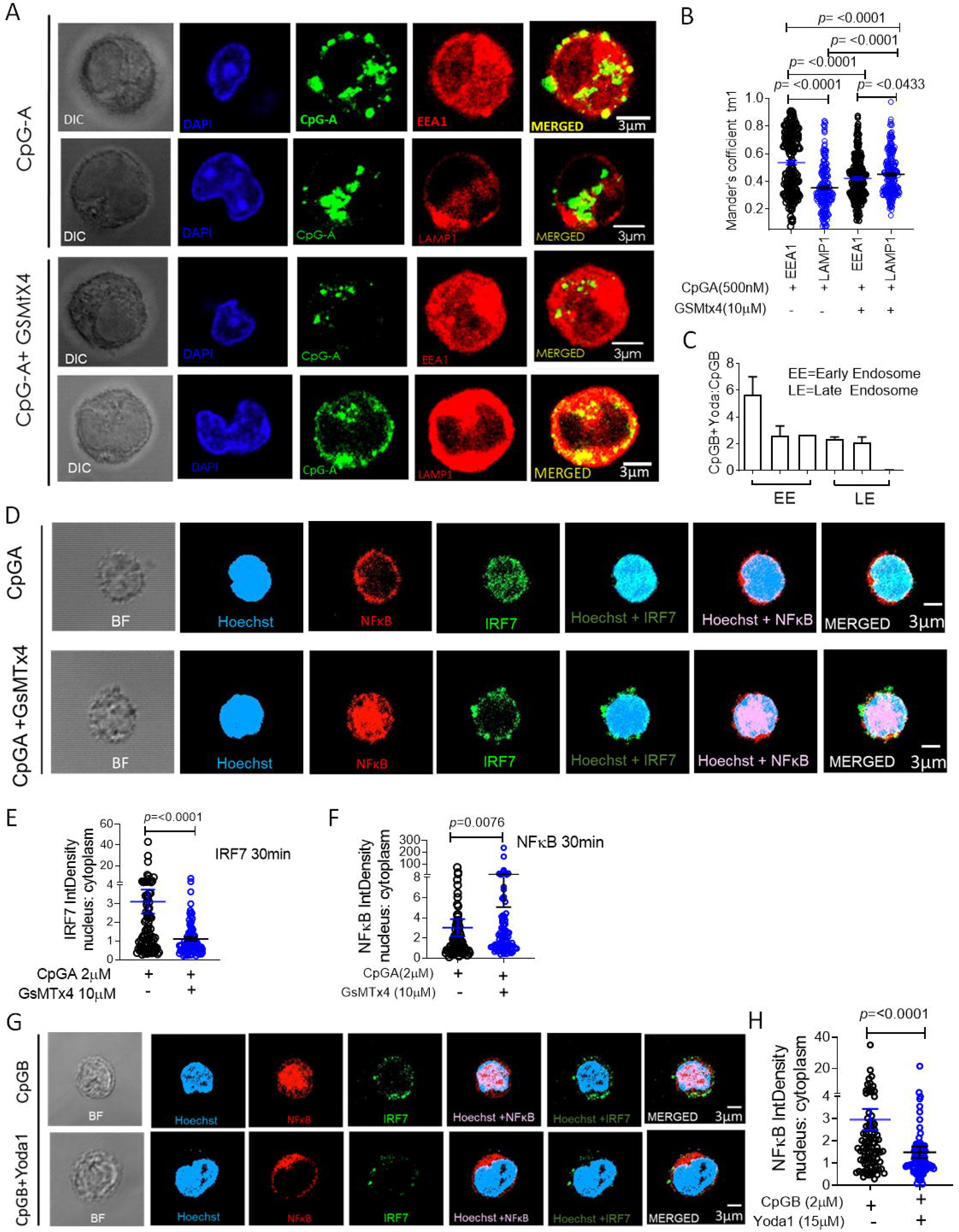
PIEZO1 activation promotes CpGA retention in early endosomes and IRF7 nuclear translocation. **A**) Representative confocal images of fixed, immunostained primary human pDCs following CpGA-FITC stimulation, visualizing early endosomes (EEA1) and late endosomes (LAMP1) under basal and GsMTx4 treated conditions. **B**) Manders’ coefficient analysis quantifying co-localization of CpGA with EEA1 and LAMP1 under control and PIEZO1 inhibited (GsMTx4 treated) conditions (n>195 cells per condition). **C)** Bar graphs showing relative detection (OD 450 nm) of biotinylated CpGB in subcellular fractions of pDCs stimulated with CpGB-biotin+/- Yoda1. Data represent mean ± s.e.m and is derived from two independent experiments. **D**) Representative confocal images of fixed, immunostained human primary pDCs showing subcellular localization (nuclear versus cytosolic) of NF-κB and IRF7 following CpGA stimulation under control and PIEZO1 inhibited conditions. **E,F**) Quantitative analyses of nuclear-to-cytoplasmic (N:C) intensity ratios of IRF7 (**E)** and NF-κB **(F)** in CpGA stimulated pDCs treated with or without GsMTx4. **G**) Representative confocal images of subcellular localization of NF-κB in primary human pDCs treated with CpGB in the presence or absence of Yoda1. **H**) Quantification of N:C intensity ratios of NF-κB in pDCs stimulated with CpGB in the presence or absence of Yoda1 (n>90 cells per condition). All data is representative of at least three independent experiments. Student’s t-test was used to calculate significance and data is represented as mean ± s.e.m. Statistical analysis was performed using Student’s t-test **(B)** or Mann-Whitney U test **(E,F, H)**

We next investigated the functional consequence of altered endosomal trafficking on downstream signaling and transcription factor localization. Nuclear localization studies using high-resolution imaging showed that CpGA retention in early endosomes is associated with robust translocation of IRF7 to the nucleus (Figure 3D, Supplementary Video 6), a key step for initiating type I IFN gene transcription (Honda et al, 2005; Honda et al, 2006). In contrast, inhibition of PIEZO1 significantly impaired nuclear localization of IRF7 following CpGA stimulation and led to increased nuclear translocation of NF-κB consistent with enhanced CpGA routing to late endosomes (Figure 3D-F, Supplementary Video 7). We also assessed whether PIEZO1 activation could alter CpGB-driven nuclear translocation of transcription factors. As expected, CpGB stimulation led to prominent NF-κB translocation (Figure 3G) into the nucleus. However, co-treatment with the PIEZO1 agonist Yoda1, attenuated NF-kB translocation to the nucleus, although significant nuclear translocation of IRF7 was not discernible at the same timepoint (Figure 3G, H). Taken together, these results demonstrated that PIEZO1 plays a key role in controlling the endosomal trafficking of CpGA, prolonging its retention in early endosomes thereby facilitating sustained IRF7 activation and IFN-α production in human pDCs.

### PIEZO1-driven calcium influx induces localized actin remodeling to stabilize CpGA-containing early endosomes in human pDCs

We next investigated the molecular mechanisms by which PIEZO1 activation promotes CpGA retention in early endosomes. PIEZO1 is a calcium channel and based on prior evidence from our group linking PIEZO1 channels to cytoskeletal regulation (Liu et al, 2018; Liu et al, 2024), we hypothesized that activation of PIEZO1 and local accumulation of Ca^2+^ may contribute to peripheral retention of the early endosomes through calcium-dependent local F-actin remodeling. To test this, first we measured intracellular calcium levels using a fluorescent calcium indicator (Fluo-3AM) and flow cytometry. Both CpGA and Yoda1 induced a robust calcium influx in primary human pDCs, which was abolished following PIEZO1 knockdown (Figure 4A, B), confirming PIEZO1 dependence. We then assessed whether PIEZO1-driven calcium signaling influences actin distribution, as documented earlier in the context of T cell activation and migration (Liu et al, 2018; Liu et al, 2024). To examine the spatial organization of the filamentous actin (F-actin) network, we performed confocal microscopy of cells co-stained for F-actin and EEA1 or F-actin and LAMP1. Images were analyzed ensuring the cortical F-actin (underlying the plasma membrane) was disregarded and only internal F-actin patches were considered (Supplementary Figure 4A, B). The data demonstrated higher F-actin enrichment around early endosomes (higher colocalization of actin-rich patches with EEA1-rich structures), compared to late endosomes (labelled with LAMP1) in CpGA treated pDCs (Figure 4C, D, Supplementary Figure 4A, B), suggesting preferential actin remodeling locally around the early endosomes. Importantly, this F-actin enrichment around early endosomes was significantly reduced upon PIEZO1 inhibition with GsMTx4, further supporting its dependence on PIEZO1-mediated signaling (Figure 4C, D).

**Figure 4:**
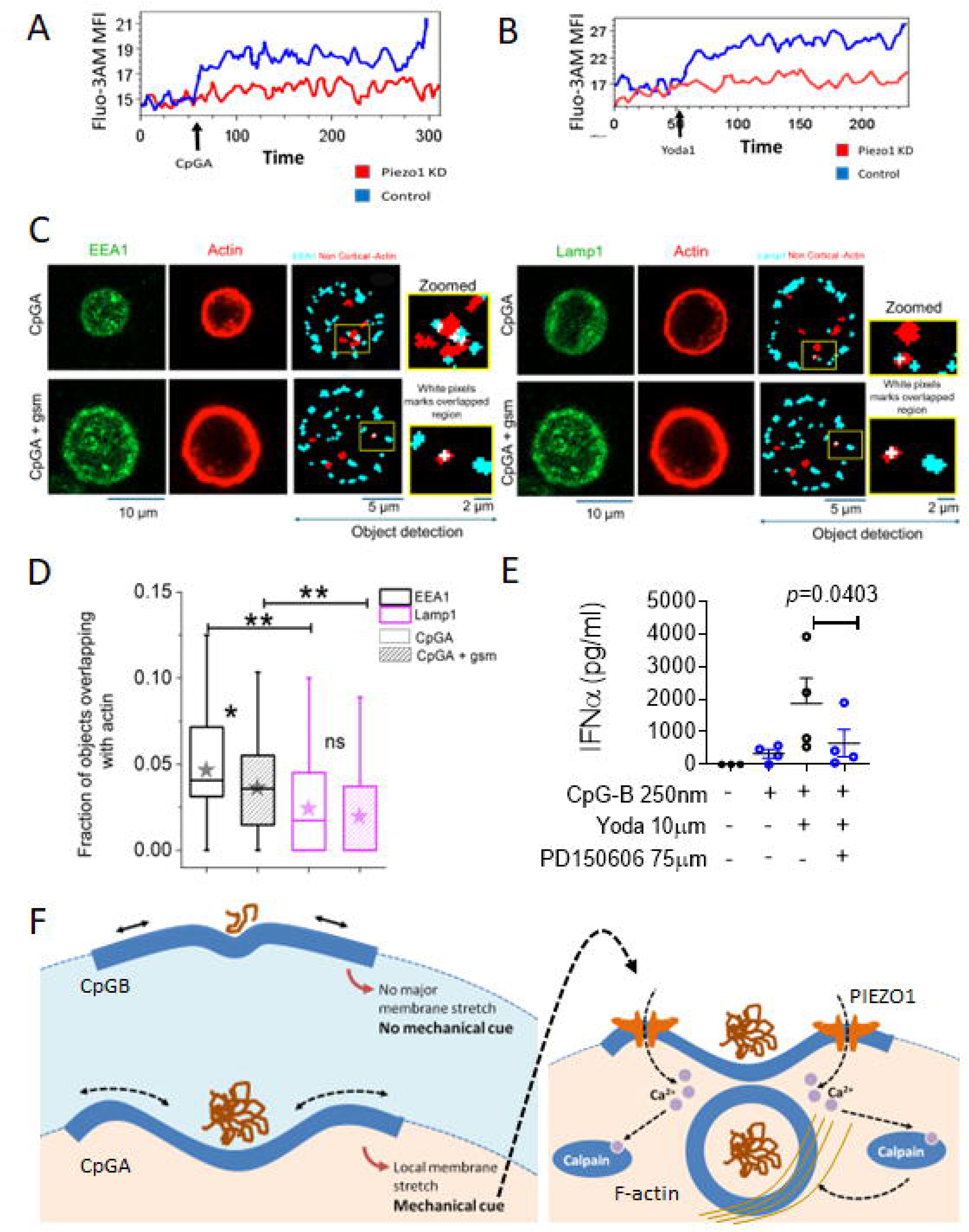
PIEZO1-driven calcium influx induces localized actin remodeling to stabilize CpGA-containing early endosomes. **A,B)** Intracellular Ca2+ abundance in control and PIEZO1 KD pDCs treated with CpGA **(A)**, or Yoda1 **(B)** was assessed by flow cytometry. Arrows represent the time-point of addition of the indicated treatments. **C)** Representative images of dual channel confocal images of EEA1/Lamp1 in the green channel and actin labeled in red channel under CpGA and CpGA +GsMTX4 treated condition - along with the typical object detection (right side panel) observed away from the cortex (without cortical actin) for a particular z-slice of a cell (insets shows the zoomed-in view of a few colocalized objects in each condition).The white pixel regions indicate the colocalized region of EEA1/Lamp1 with non-cortical actin. xy-1 pixel = 0.13 µm, z step – 1 µm **D)** Cell -wise comparative study of the fraction of endosomal objects that overlap with actin under a particular treatment with ** indicating a significant decrease in actin colocalization with early endosomes on PIEZO1 inhibition (without cortical actin). EEA1 (N_cell_): CpGA – 116, CpGA + gsm – 116; Lamp1 (N_cell_): CpGA – 136, CpGA + gsm – 108. Scale bar-10 µm. Mann–Whitney U statistical significance test where ns-p >0.05, * - p <0.05, ** - p <0.001. **E)** IFNα production in supernatants of pDCs stimulated with CpGB in the presence of indicated treatments. n=4. Student’s T-test was performed. Data is representative of two independent experiments and represents mean+/-s.e.m.

Next, to test whether F-actin remodeling is essential for PIEZO1-driven type I IFN production, we pharmacologically disrupted F-actin polymerization using the calpain inhibitor PD150606. This significantly reduced IFN-α production in cells co-stimulated with CpGB and Yoda1 (Figure 4E). Cumulatively, these findings demonstrated that PIEZO1-dependent calcium influx initiates a localized actin remodeling program that stabilizes CpGA-containing early endosomes, thereby sustaining IRF7 activation and robust type I IFN responses in human pDCs (Figure 4F).

## Discussion

The present study uncovers a previously unrecognized role for PIEZO1-mediated membrane tension sensing in regulating nucleic acid-induced type I interferon production in pDCs (Fig. 4F), revealing that innate immune responses are critically shaped not just by ligand chemistry or receptor specificity, but also by the biophysical properties such as ligand’s propensity for self-association increasing the cargo size and membrane tension. We demonstrate that in physiological condition CpGA molecules, unlike CpGB, self-associates to form large aggregates that impose significant membrane stretch during uptake. This tension activates PIEZO1 at the plasma membrane, triggering calcium influx and localized actin polymerization that stabilizes CpGA within early endosomes longer permitting sustained IRF7 activation and robust type I IFN production. Inhibition of PIEZO1 or downstream actin filament polymerization leads to premature CpGA trafficking away from early endosomes and abrogates interferon responses. Strikingly, chemical activation of PIEZO1 upregulates IFN-α production in response to CpGB, underscoring the sufficiency of mechanotransduction in rewiring pDC signaling.

While much is known about pattern recognition receptor (PRR) specificity and endosomal trafficking in shaping innate immune signaling (Brubaker et al, 2015), this study introduces membrane tension and mechanosensing as critical upstream regulators of these pathways. The observation that CpGA-induced membrane deformation activates PIEZO1 to promote actin remodeling and endosomal retention positions PIEZO1 as a mechanosensory gatekeeper between ligand uptake and compartmentalized signaling. Notably, previous studies have described the importance of early endosomal localization for CpGA-mediated IRF7 activation (Honda et al, 2005; Guiducci et al, 2006), but the cues regulating this phenomenon remained elusive. Our findings provide a mechanistic link between physical properties of the ligand and subcellular signal routing, offering a new layer of regulation in the context of human pDC biology.

These insights also expand the current understanding of innate immunity beyond chemical recognition. Increasing evidence supports that physical attributes such as size, shape, and rigidity influence immune responses (Benne et al, 2016; Baranov et al, 2021). We now show that size of the CpGA cargo, through mechanical activation of PIEZO1, modulates TLR9 signaling. This may explain why CpG-based immunotherapies optimized solely for sequence or chemical stability have shown limited efficacy (Chen et al, 2021). Our findings suggest that tuning the biophysical features of nucleic acid ligands could offer a new strategy to modulate interferon responses. While PIEZO1 has been implicated in other immune cell functions (Liu et al, 2018; Atcha et al, 2021; Liu et al, 2024), this is the first report linking PIEZO1 to nucleic acid sensing and regulation of type I IFN induction by human pDCs.

Given the central role of pDCs and type I IFNs in autoimmune diseases such as lupus (Ganguly et al, 2013; Ganguly, 2018), our findings may hold particular relevance in these contexts. In autoimmune diseases, self-nucleic acids frequently form large complexes with the antimicrobial peptide LL37, which is an established danger molecule in different autoimmune contexts, viz. psoriasis and systemic lupus (Lande et al, 2007; Ganguly et al, 2009), or with autoantibodies (Barbalat et al, 2011; Theofilopoulos et al, 2017). These complexes also undergo aberrant endo-lysosomal sorting, leading to their accumulation triggering excessive type I IFN production, a key driver of inflammation and pathology (Lande et al, 2007; Ganguly et al, 2009). Our findings can be extrapolated to suggest that such endogenous immune complexes carrying TLR ligands may lead to increase in local membrane tension during their uptake by pDCs, potentially activating PIEZO1 and modulating downstream type I IFN induction. These insights also identify PIEZO1 as a potential therapeutic target for modulating dysregulated IFN responses in the contexts of autoimmune diseases and autoreactive inflammation wherein a crucial role of pDC-derived type I IFNs are established (Ganguly, 2018).

The present study also raises compelling new questions. Is PIEZO1 a general modulator of nucleic acid sensing across other endosomal PRRs such as TLR7, TLR8, or TLR3? And might endogenous nucleic acid containing immunogenic complexes such as those observed in necrotic tissue or neutrophil extracellular traps exploit this pathway to drive autoimmune inflammation? These avenues of inquiry could extend the impact of our findings beyond CpG biology and into fundamental mechanisms of self-nonself discrimination and immune homeostasis.

Interestingly, a recent study also showed that chronic extracellular matrix stiffening in fibrotic skin suppresses TLR9-induced type I IFN production in pDCs via NRF2 activation, emphasizing the role of sustained mechanical cues in pDC regulation (Chaudhury et al, 2025). In contrast, our findings highlight that acute, ligand-intrinsic membrane tension driven by uptake of larger CpGA cargo activates Piezo1 to enhance IFN responses. These differing outcomes likely reflect distinct magnitudes, durations, and cellular/clinical contexts of mechanical input. Together, the findings suggest that pDCs integrate diverse mechanical signals to modulate type I IFN output in a context-dependent manner.

In summary, our work reveals that Piezo1 functions as a biophysical checkpoint that translates membrane tension into durable type I IFN signaling in pDCs, resolving a key mechanistic gap in TLR9 signaling biology and redefining how physical properties of ligands shape innate immune responses. By identifying mechanosensation as a central determinant of nucleic acid–induced immunity, we introduce a new conceptual axis in pDC regulation, one that integrates mechanical force, membrane dynamics, cytoskeletal remodeling, and transcriptional activation, and opens new therapeutic possibilities for harnessing or modulating interferon responses in health and disease with wide-ranging implications for autoimmunity, vaccine design, and immunotherapy.

## Methods

### Small angle X-ray scattering

The SAXS datasets for studying the changes induced by solvent constitution on CpGA (ODN2216, InvivoGen) and CpGB (ODN2006, InvivoGen) were acquired on SAXSpace instrument (Anton Paar GmbH, Graz, Austria at CSIR-Institute of Microbial Technology, Chandigarh, India). Line collimation arising from a sealed X-ray tube was incident on the samples and buffer contained in a thermostated quartz capillary. For each dataset, one frame of one hour of exposure to X-rays was considered. The scattering data was recorded on a 1D CMOS Mythen detector (Dectris, Switzerland). Finally, the scattering data profile was collected as I(s) as a function of s, where s is momentum transfer vector defined as s = 4π(sinθ/λ) and units in 1/nm. The intensity profiles were collected for both CpGA and CpGB in water and PBS, and blank solvents from 5 to 50°C at an interval of 5-10°C. Supplementary Table S1 summarizes all the programs used to process the acquired data. After the data collection, the beam position was set to zero using the software SAXSTreat and SAXSQuant was used for buffer subtraction and desmearing the data to represent true point collimation. The SAXS data profiles of both the samples, at different temperatures were analyzed to obtain radius of gyration (R_g_), maximum linear dimension (D_max_) and distance distribution function of pairwise vectors (P(r)) using the ATSAS 3.0.3 version. Using Guinier approximation for globular or rod-like scattering shape profiles provided R_g_ and radius of cross-section (R_c_), respectively. Persistence length of the molecules, L and their aspect ratio, A were estimated using L = (12((R_g_^2^ – R_c_^2^)))^1/2^ and A = R_g_/R_c_, respectively. Kratky analysis was also done using the same program. All the plots were made using ATSAS PrimusQT suite of programs. Shape restoration was done using ATSAS online server. Briefly, for all sets, 10 runs of DAMMIF program were done to model dummy residue uniform density models without any shape or symmetry bias. The solutions were aligned and averaged using offline version of DAMAVER suite of programs. Molecular maps were generated for the average and variable of the ten dummy residue models. The images of the models were made using UCSF Chimera program version 1.13.

### Fluorescence correlation spectroscopy

The self-association of the CpGA and CpGB were measured using Fluorescence Correlation Spectroscopy (FCS). FCS analyzes fluctuations in fluorescence intensity within the confocal volume to determine the diffusion times of fluorescent molecules present in nanomolar concentration. 20nM Alexa-488 labeled CpGA in the presence of unlabeled CpGA concentration ranging from 0-5µM was used. Similarly, 20nM Alexa-488 labeled CpGB in the presence of unlabeled CpGB concentration ranging from 0-5µM was used. The FCS measurement was performed using an ISS Alba FFS/FLIM confocal system (Champaign, IL, USA), integrated with a Nikon Ti2U microscope (Nikon, Japan) equipped with the Nikon CFI PlanApo 60X/ 1.2NA water immersion objective. The samples were drop casted on the 22 mm grease-free coverglass (Blue-Star, India). The samples were excited one at a time using a 488-nm picosecond pulsed diode laser and FCS measurements were acquired. The fluorescence emission was detected using a SPAD (Single Photon Avalanche Detector) detector and the 530/43-nm band-pass filter. The FCS correlation curves obtained were fit to the 3D Gaussian 1-component diffusion model using VistaVision software (ISS Inc., USA). The fitted FCS correlation curves were plotted using OriginPro24 software (OriginLab, USA).

In the 3D Gaussian diffusion model, which considers a single type of diffusing molecule (the 3D Gaussian 1-component model, excluding triplet state contributions), the correlation function G(τ) is defined by the following equation:

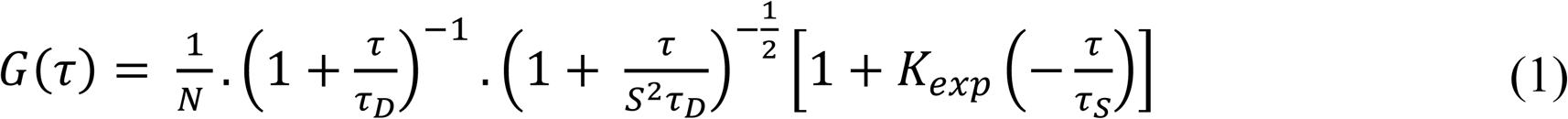

where *τ*_*D*_ represents the diffusion time of the molecules, N is the average number of molecules within the observation volume, and S is the structural parameter that defines the ratio between the radius and the height of the confocal volume. The value of *τ*_*D*_, obtained by fitting the correlation function, is related to the diffusion coefficient (D) of a molecule through the following equation:

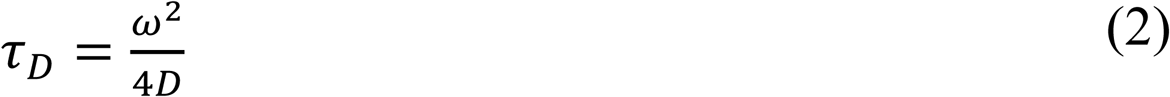

where ω is the size of the observation volume. The beam waists in the radial and axial dimensions were calibrated using a standard fluorescence dye, Rhodamine 6G (R6G), in water with a known diffusion coefficient of 2.8 × 10⁻⁶ cm²/s. The fold change in the diffusion time was calculated with the mean diffusion time of 20nM CpGA. The Number of the particle (N) in the confocal volume was calculated using VistaVision software for the respective species.

### Isolation and culture of human plasmacytoid dendritic cells

Peripheral blood samples were obtained from healthy adult donors after informed consent, the approval from institutional Human Ethics Committee at CSIR-Indian Institute of Chemical Biology, Kolkata, India, in accordance with the Declaration of Helsinki. Buffy coats were collected from the Blood Ban of Tata Medical Center, Kolkata, India, under an approved material transfer agreement. Blood was diluted 1:2 with PBS and layered over a Hisep LSM gradient for density centrifugation (2,500 rpm, 20 min, 25 °C, brake and acceleration settings = 1). The mononuclear cell layer was collected, washed, and subjected to hypotonic red blood cell lysis. Peripheral blood mononuclear cells (PBMCs) were resuspended in MACS buffer (PBS containing 0.5 mM EDTA and 0.5% BSA), and pDCs were isolated by magnetic-activated cell sorting (MACS) using anti-BDCA4–coated magnetic beads (Miltenyi Biotec) following Fc receptor blockade. Labeled cells were passed through LS columns on a magnetic stand, washed thoroughly, and eluted to obtain highly purified pDCs. Cells were cultured in complete RPMI 1640 medium in 96-well U-bottom plates at 37 °C with 5% CO₂ and processed as described in the figure legends.

### RNA interference in human pDCs

Primary human pDCs were subjected to siRNA-mediated knockdown using the 4D-Nucleofector™ System (Lonza) according to the manufacturer’s instructions. Briefly, cells were resuspended in 100 µl of supplemented P3 Primary Cell Solution and nucleofected with 175 ng of either a non-targeting EGFP control siRNA or a Piezo1-specific siRNA or a ARHGAP26 siRNA using the program FF168. Following nucleofection, cells were immediately transferred to RPMI 1640 medium supplemented with 10% fetal bovine serum and incubated at 37 °C in 5% CO₂ for 18 hours prior to downstream analyses. Knockdown efficiency was validated by quantitative PCR to assess *ARHGAP26* and *PIEZO1* mRNA levels.

### Flow cytometric analysis of CpG oligonucleotide uptake by human pDCs

CpGA and CpGB oligonucleotides were fluorescently labeled using either Alexa Fluor 647 or Alexa Fluor 488 dyes, following the protocol provided with the Ulysis Nucleic Acid Labeling Kit (Life Technologies). Primary human pDCs, pre-incubated for 20 mins with Pitstop2 or Genistein before they were incubated with fluorescently tagged CpGA or CpGB for 30 minutes at 37 °C. Following incubation, cells were washed thoroughly with PBS to remove unbound material and immediately analyzed on a flow cytometer (BD LSRFortessa) to assess internalization based on fluorescence intensity.

### Scanning electron microscopy

Samples were prepared for scanning electron microscopy according to a previously published protocol (Langthasa et al, 2021). Briefly, primary human pDCs were seeded onto poly-L-lysine– coated glass coverslips and allowed to adhere for 4 hours at 37 °C. Cells were then stimulated with CpGA or CpGB for 10 minutes, followed by fixation in 2.5% glutaraldehyde at 4 °C overnight. After fixation, samples were washed three times with phosphate-buffered saline (PBS; 5 minutes each) to remove residual fixative, followed by five rinses with distilled water to eliminate salt residues. Cells were dehydrated through a graded ethanol series (30%, 50%, 70%, 90%, and 100%), and coverslips were air-dried completely at room temperature. Samples were imaged using an Environmental Scanning Electron Microscope (ESEM Quanta, Thermo Fisher Scientific).

### Interference Reflection Microscopy (IRM): Global Tension measurement

An inverted microscope (Nikon, Japan) with 60x 1.2 NA water-immersion objective, Hg arc lamp and necessary interference filter (546 ± 12 nm) were used for the interference reflection microscopy (Biswas et al, 2017; Limozin et al, 2009). IRM imaging of pDCs was achieved by using magnetic beads and magnet under the coverslip to bring them closer to the glass substrate and enhance interaction with the substrate. Cells that were significantly adhered were used for imaging. Fast time-lapse images at an interval of 20 ms with 50 frames per second frame rate were captured using a sCMOS camera over 8192 frames was done to capture the fast transient changes in mechanical properties caused by cargo oligo-nucleotide addition (Hamamatsu, Japan).

### Total Internal Reflection Fluorescence (TIRF) microscopy coupled with IRM microscopy: Local Tension measurement

The cells were kept on the microscope stage top incubator (Tokai Hit, Japan) for 10 min for the system to stabilize. For correlative TIRF-IRM sequential imaging, Eclipse Ti-E motorized inverted microscope (Nikon, Tokyo, Japan) equipped with 100x - 1.4 NA water-immersion objective and sCMOS (ORCA Flash 4.0 Hamamatsu, Japan) was used. For IRM-based imaging a 100W Hg arc lamp with necessary interference filters (546 ± 12 nm),50-50 beam splitter was used (Biswas et al, 2017; Biswas et al, 2019; Chakraborty et al, 2022). For TIRF imaging coherent OBIS laser of 488 nm was used (Fish, 2009; Poulter et al, 2015; Fish, 2022).

Human pDCs cells were seeded on poly D-lysine coated confocal dishes and were incubated at 37C, 5% CO_2_ for 45 min for them to adhere to the glass substrate. After 45 min of cell seeding the media was replenished and cells were taken for imaging. Properly adhered cells were chosen for imaging. Calibration for membrane fluctuation tension mapping was done using 60µm diameter polystyrene NIST beads (Bangs Laboratories Inc.). The sequence followed for correlative TIRF-IRM sequential imaging for any field that had fluorescent signal was TIRF-IRM-TIRF. For time-points before oligonucleotide addition (no fluorescence), only IRM frames were captured. For IRM imaging, time-lapse images at an interval of 50 ms with 20 frames per second rate were captured. For a particular field 2048 frames were captured for effective membrane fluctuation tension measurement. This first IRM movie was considered as the baseline without the oligo-nucleotide addition. 1µM of FITC-tagged CpGA was subsequently added to the media without disturbing the field of view. Following the same field at 5 min, 15 min and 20 min of addition of oligo-nucleotides sequential TIRF-IRM images were captured. For TIRF-Imaging, images were acquired at an interval of 200 ms for 300 of the same field as that of IRM at a frame rate of 3.33 frames per second with a penetration depth of approximately 90-100 nm. TIRF movies captured before and after the IRM movie were used to verify the stability of the observed fluorescent pattern over the 1-2 min imaging period. CpGB-FITC being a very small tagged oligonucleotide, obtaining properly adhered pDCs, fluorescently labelled with CpGB cargo was not feasible using TIRF-IRM system.

### Analysis of IRM data

Proper adherence of the cells to the glass substrate was ensured for selecting them for analysis. Further, from these selected adhered cells only regions that were within a range of ∼100 nm from the coverslip – also called the first branch region (FBR) were chosen and the size of these chosen FBRs were 4 × 4 pixels corresponding to ∼ 65 nm x 65 nm per pixel for IRM images of the TIRF-IRM sets and ∼ (300 nm)^2^ for only IRM images. To maintain stringency in selecting the subset of regions for extracting parameters and estimating the membrane height fluctuations, the methodology designed previously (Biswas et al, 2017; Biswas et al, 2019) has been followed.

For analyzing the mechanical parameters of the cells under these two conditions over the time points, the power spectral density (PSD) was estimated by autoregression technique (probability of covariance or pcov) (MATLAB, Mathworks Inc, USA). For estimating active temperature (A), effective cytoplasmic viscosity (η_eff_), confinement (γ) and membrane tension (σ), PSD is fitted to the Helfrich-based Theoretical Model(26:

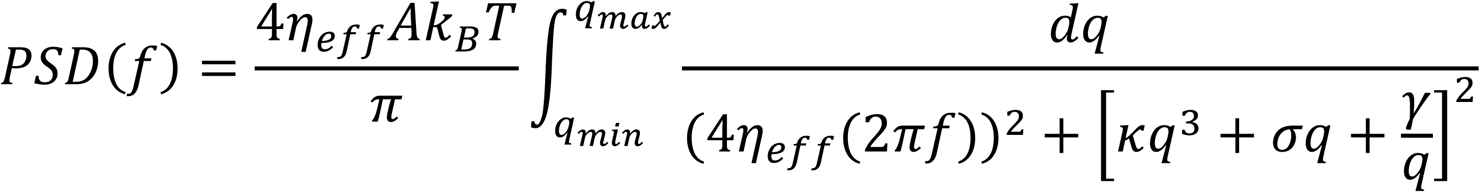

The fitting was done using MATLAB and fits with *R*^2^ > 0.9 were considered (Biswas et al, 2019). The standard deviation of the membrane height over time series of a single pixel is estimated and averaged over single FBRs to obtain the membrane height fluctuations termed as SD_time_. Generally, effective membrane tension is termed as the fluctuation tension as the interpretation is dependent on the framework of membrane fluctuations (Shiba et al, 2016). The effective membrane fluctuation tension is termed as ‘tension’ in the text. It has contributions from all players affecting membrane fluctuations, including the cytoskeleton.

For tension mapping (Figure1G), PSDs calculated from each pixel were fitted and all parameters extracted from fits—including tension and *R*^2^. The pixel-wise standard deviation of the membrane height (SD_time_) over time series was also mapped.

### TIRF-IRM correlative analysis

The first frame of the TIRF movie obtained after the IRM movie was used to obtain the pixel-wise correlation of local intensity with pixel-wise-fitted tension and pixel-wise SD_time_ at the sites where FITC-tagged oligonucleotide binds to plasma-membrane. The IRM movies for every time points were template matched (using Fiji ImageJ software) with respect to time points as well as the fluorescence fields. For every cell at a particular time point after oligonucleotide addition. At the peak of fluorescence of each cluster in cells in the first frame of the TIRF movie obtained after the IRM movie, line scans (11 pixels along x and 11 pixels along y direction) are performed for 10-20 clusters of CpGA per time point. These line scans were mapped onto pixel-wise tension and pixel-wise SD_time_. This correlative mapping was carried out for the baseline (no treatment) as well as 5 min, 15 min and 20 min after addition of oligonucleotides. For all the time points the first frame of TIRF movie captured after its corresponding IRM movie was used for this correlative mapping. For plotting the baseline condition, the regions where clusters appeared on the first frame of the first TIRF movie captured after 5 min of addition CpGA were used to trace back the basal local membrane height fluctuation and tension in the untreated condition at those local sites where the oligonucleotide binds after addition. The center is normalized to one for all scans of CpGA/CpGB clusters followed by averaging over 10-20 clusters per time point. The same cells were followed in time, but puncta were independently chosen for each time point. Error is denoted as SEM. The profiles were also averaged based on distance from center. Only for better representation purposes both negative and positive distance were plotted – however the data is same for two points at same distance from the center. CpGA/CpGB puncta very close to cell edge were not taken into account to avoid arbitrary tension values expected for pixels lying outside the cell. During the correlation mapping at the clusters, pixels where PSD fitted to *R*^2^ > 0.9, membrane fluctuation tension <5000 pN/µm and SD_time_ < 15 nm were only considered. Mann–Whitney U test was performed to calculate the statistical significance of the various mechanical properties at various time points with respect to the baseline. From the center of the cluster, the increase in tension within 325 nm was calculated and the probability distribution of maximum surge in tension was plotted for all the conditions. Also, the mean pixel-wise local tension for 10X10 pixels around the center of the cluster, normalized with respect to baseline condition, was also calculated and plotted.

### Immunostaining for confocal imaging

Primary human pDCs were seeded onto poly-D-lysine coated (1 µg/ml) coverslips at a seeding density of 0.05 × 10^6^ cells per coverslip and incubated at 37 °C in 5% CO_2_ for 2-3 hours for attachment. For PIEZO1 localization studies, cells were either left untreated (control) or stimulated with either CpGA (250nm) or CpGB (250 nm). Post incubation, cells were fixed in 4% paraformaldehyde for 15 min at room temperature. Following fixation, cells were blocked in 3% bovine serum albumin in PBS containing 0.2% Triton X-100 for 60 minutes at room temperature. Cells were incubated overnight at 4°C with rabbit anti-human Piezo1 antibody (proteintech), at 1:100 dilution (5 µg/ml). Next day, Alexa 568-conjugated anti-rabbit IgG (1:200 dilution) was applied for 60 minutes at room temperature. After washing, coverslips were mounted with Vectashield containing DAPI for nuclear staining and sealed for confocal imaging.

For, endosomal colocalization analysis, pDCs were pretreated with GsMTX4 (10µM) for 30 minutes, followed by addition of FITC tagged CpGA for 15 minutes. Cells were washed with PBS to remove unbound CpGA and seeded onto precoated poly-D-lysine (1µg /ml) coated coverslips for adherence. After 15minutes, cells were fixed and blocked as described above. Immunostaining was performed using mouse anti-human EEA1 (eBioscience) and rat anti-human LAMP1(Abcam), both at dilution of 1:100 (5 µg/ml) followed by overnight incubation at 4°C. Alexa 568-conjugated anti-mouse IgG & Alexa 647-conjugated anti-rat IgG secondary antibodies were applied at 1:200 dilution for 60 minutes. Cells were mounted with Vectashield containing DAPI prior to confocal imaging.

For assessment of transcription factor nuclear transocation, primary human pDCs were stimulated for 30 minutes with 2µM CpGA in the presence or absence of GsMTx4 (10µM) or stimulated with CpGB in the presence or absence of Yoda1 (15µM). Cells were fixed, permeabilized and blocked as described above. Immunostaining was performed using mouse anti-human phospho-IRF7 (Ray Biotech 128-10356-3) and rabbit anti-human NF-κB-p65 antibodies (both at 1:100 dilution). Subsequent detection was performed using goat anti-rabbit Alexa Fluor 568 and goat anti-mouse Alexa Fluor 488 antibodies (both at 1:200 dilution). Nuclei were counterstained with Hoechst dye. For quantification of nuclear versus cytoplasmic fluorescence intensity density, regions of interest (ROIs) were generated in ImageJ/Fiji as follows: nuclei were segmented using the DAPI channel—to create a clean nuclear mask (ROIₙ) The full cell boundary was delineated using bright-field images, with manual ROI tracing to produce a whole-cell ROI (ROIₓ). The cytoplasmic ROI (ROI_c_) was then defined by subtracting the nuclear mask from the whole-cell ROI (ROI_c_ = ROIₓ – ROIₙ). Fluorescence intensity density of NF-κB and IRF-7 was measured separately within ROIₙ (nuclear) and ROI_c_ (cytoplasmic) regions. Finally, the nuclear-to-cytoplasmic intensity ratio was calculated by dividing the nuclear intensity density by the cytoplasmic intensity density.

For endosomal and actin co-localization studies, the protocol for endosomal staining was followed. For actin visualization primary human pDCs were stained with Alexa flour 532 tagged phalloidin (1:200 dilution; 0.5µg/ml) following CpGA addition in the presence or absence of GsMTx4.

### Confocal microscopy image Analysis

For counting the fraction of compartments endosomal or puncta that have spatial co-distribution with actin puncta (in MATLAB), a mask was applied over the image to select the cell. For this cell, the equatorial z slices were selected, and thresholding was performed using the appropriate threshold, resulting in a binary image for both the endosomal and actin signal. Single pixels were removed using serial erosion and dilation, and subsequently, the binary image was used to detect objects. It was observed that the actin objects from the cortex were detected as a single large continuous object. So, for selecting actin object away from the cortex, apart from choosing the threshold based on the local mean intensity (first-order statistics) in the neighbourhood of each pixel and giving a sensitivity factor towards thresholding more pixels as foreground, where more the sensitivity more pixels are considered as foreground, an object-size-based threshold was also applied. Objects were detected for each channel separately. The pixels overlapping in the two binary images were considered to be colocalizing and another binary image/matrix for the colocalized pixels were constructed. Colocalized objects were detected from this new formed-matrix using 8-pt connectivity (MATLAB) object detection. The sum of number of all the colocalized objects in all the z-slices divided by the total number of endosomal objects detected in all z-slices gave us the fraction of objects that overlapped with actin per cell. Mann-Whitney U test was used to calculate significance, unless specified. (ns denotes p>0.05, p<0.05 is denotes as * and p<0.001 is denoted as ** statistical significance).

### ELISA for IFN-α

Concentration of IFN-α in pDC culture supernatants was determined using sandwich ELISA (Mabtech, Sweden) according to manufacturer’s protocol.

### RNA isolation and real time PCR

Total RNA was extracted from using TRIzol reagent (Invitrogen), following the manufacturer’s protocol. RNA was reverse-transcribed into complementary DNA (cDNA) using the SuperScript III First-Strand Synthesis System (Invitrogen). Quantitative real-time PCR was performed on an Applied Biosystems 7500 Fast Real-Time PCR System using SYBR Green chemistry. Primer sequences used for gene expression analysis are provided in Supplementary Table S2.

### Flow cytometry for calcium influx

Isolated primary human pDCs were stained with the calcium-sensitive fluorescent dye Fluo-3 AM (1.5 µM; Invitrogen) for 30 minutes at 37 °C in PBS supplemented with 1.2 mM CaCl₂ and 2% fetal bovine serum (GIBCO). After incubation, cells were washed twice in the same buffer and incubated at room temperature for an additional 30 minutes to allow complete de-esterification and efflux of unincorporated dye. Stained cells were then acquired on a BD LSRFortessa™ flow cytometer at the indicated time points before and after the addition of specific treatments. Changes in intracellular calcium levels were monitored by measuring the mean fluorescence intensity (MFI) in the FITC channel (corresponding to Fluo-3 AM), with increased MFI reflecting calcium mobilization in response to stimulation.

### Subcellular fractionation

Subcellular fractionation was performed according to a previously published protocol (Bose et al, 2017). Primary human pDCs were incubated with biotinylated CpGB (CpGB-biotin, 5 µM) in the presence or absence of Yoda1 (10 µM) for 30 minutes at 37 °C. After stimulation, cells were immediately transferred to ice, washed with cold PBS, and processed for subcellular fractionation using an iodixanol gradient. Briefly, 2-3 × 10^6^ cells per condition were resuspended in hypotonic lysis buffer containing 50 mM HEPES (pH 7.8), 78 mM KCl, 4 mM MgCl₂, 8.4 mM CaCl₂, 10 mM EGTA, 250 mM sucrose, 100 μg/ml cycloheximide, 5 mM vanadyl ribonucleoside complex, and an EDTA-free protease inhibitor cocktail (Roche). Cells were allowed to swell on ice for 10 minutes and lysed by 20–25 strokes with a glass Dounce homogenizer (Sartorius). The homogenate was centrifuged at 1,000 × g for 10 minutes at 4 °C to remove unbroken cells and nuclei. The post-nuclear supernatant was layered onto a pre-formed discontinuous iodixanol gradient (3%–30%, prepared in gradient buffer) and ultracentrifuged at 36,000 rpm for 5 hours at 4 °C in an SW60 Ti rotor (Beckman Coulter). Fractions were manually collected from the top in equal volumes and maintained on ice. Endosome-enriched fractions were identified based on density and lysed by adding saponin to a final concentration of 0.1% (w/v), followed by incubation on ice for 20 minutes. To quantify the distribution of CpGB-biotin across fractions, 200ul of fractions were incubated overnight at 4 °C in high-binding 96-well ELISA plates (Nunc). After washing with PBS containing 0.05% Tween-20 (PBST), wells were incubated with streptavidin–horseradish peroxidase (HRP; 1:5,000) for 2 hours at room temperature. Plates were washed again with PBST, developed using TMB substrate, and the reaction was stopped with 1N HCl. Absorbance was measured at 450 nm using a microplate reader. Optical density values were used to assess subcellular distribution of CpGB under different treatment conditions.

### Statistics

Data were plotted and analyzed using GraphPad Prism v8.4.2. Statistical significance was assessed using unpaired Student’s t-tests, unless otherwise specified in the figure legends. A p-value < 0.05 was considered statistically significant. The number of independent replicates for each experiment is indicated in the corresponding figure legends.

## Acknowledgements

The study was funded by the Council of Scientific and Industrial Research, India (Grant no. FBR MLP-140) to D.G. S.P. is supported by a Senior Research Fellowship from University Grants Commission, India. D.G. was also supported by the Swarnajayanti Fellowship from the Department of Science and Technology, Government of India. B.S. acknowledges support from Wellcome Trust/DBT India Alliance fellowship (grant number IA/I/13/1/500885), SERB (grant number SERB_CRG_2336) and IISERK for the TIRF-IRM microscope. U.M. thanks IISERK for providing her fellowship.

## Author contributions

**Shrestha Pattanayak:** Investigation; Visualization; Methodology; Writing—review and editing. **Deblina Raychaudhuri:** Data curation; Formal analysis; Investigation; Visualization; Methodology; Writing—original draft; Writing—review and editing. **Purbita Bandopadhyay:** Investigation; Visualization; Methodology. **Upasana Mukhopadhyay:** Investigation; Visualization; Methodology; Writing—review and editing. **Tithi Mandal:** Investigation; Visualization; Methodology. **Chinky Shiu Chen Liu:** Investigation; Methodology. **Nidhi Kalidas:** Investigation; Visualization; Methodology. **Sumangal Roychowdhury:** Investigation; Visualization; Methodology; Writing—review and editing. **Krishnananda Chattopadhyay:** Formal analysis; Writing—review and editing. **Ashish:** Investigation; Visualization; Methodology; Formal analysis; Writing—review and editing. **Bidisha Sinha:** Resources; Supervision; Data curation; Investigation; Visualization; Methodology; Formal analysis; Writing—review and editing **Dipyaman Ganguly:** Conceptualization; Resources; Supervision; Data curation; Formal analysis; Funding acquisition; Visualization; Methodology; Writing—original draft; Project administration; Writing—review and editing.

