## Supplementary materials for "Spatio-temporal regulation of ligand trafficking and TLR9 activation involves PIEZO1 mechanosensing in human plasmacytoid dendritic cells"

Pattanayak et al.

**Supplementary table S1 & S2**

**Supplementary figures 1-4 with legends**

**Legends for Supplementary videos 1-7**

**Supplementary Table S1**

Details of instrumentation, and programs used for SAXS processing are tabulated below.

| **Instrument** | **SAXSpace (Anton Paar, Austria)** |
|---|---|
| Collimation | Line Collimation |
| Source and λ | X-rays, CuKα, 0.15414 nm |
| Detector | 1D Mythen |
| Sample to detector distance | 317.06 mm |
| Exposure time & Repeats | One Exposure of 60 minutes for samples & buffer |
| Subtraction from Solutions | Matched Buffer |
| **Programs** | |
| Data collection & Optics Control | SAXSDrive |
| Beam Position Correction | SAXSTreat |
| Buffer Subtraction & Desmearing | SAXSQuant |
| SAXS Intensity File Analysis | ATSAS Suite of Programs v 3.0.3 |
| Guinier Analysis | SAS Data Analysis |
| Distance Distribution Function | SAS Data Analysis |
| Molecular Weight Analysis | Bayesian Inference |
| Shape restoration | DAMMIF online version (ten times) |
| Averaging | DAMAVER offline version |
| Graphical representations | UCSF Chimera v 1.13 |
| Molecular maps | Molmap program integrated with Chimera |

**Supplementary Table S2**

| GENE NAME | PRIMER SEQUENCE | |
|---|---|---|
| Human βactin | 5'-GACGACATGGAGAAAATCTG3' | 5-'ATGATCTGGGTCATCTTCTC-3' |
| Human 18S | 5′-GTAACCCGTTGAACCCCATT-3′ | 5′-CCATCCAATCGGTAGTAGCG-3′ |
| Human IFNA1 | 5′-TCTACGATGGCCTCGCCCTT-3' | 5′ GGCTCGAGCCTTCTGG AACTGG-3' |
| Human Piezo1 | 5'-CCCAAGTGGAGCTCAGGCCC-3' | 5'-GGGCCAGGGACAGGCAGAAG-3' |

#### Supplementary Figure Legends:

**Supplementary Figure 1: A, B)**: Upper panels show the SAXS datasets acquired from samples of CpGA and CpGB molecules dissolved in water and PBS at different temperatures. Black and blue ellipses represent the average SAXS profile for CpGA and CpGB molecules in their respective plots. Inset in the panels show the Guinier approximation for globular profile and red line is the linear fit to the approximation. Mid panels show the dimensionless Kratky plots for the average datasets and insets show residuals of the respective Guinier approximation. Lower panels show the computed P(r) curves for the average datasets for CpGA and CpGB molecules. Insets in latter panels show the approximated profile (red line) in the SAXS data used to calculate P(r) curves. **C)** Molecular maps of the envelope shapes calculated for the CpGA and CpGB molecules in water are presented which doesn't show any significant difference between the two oligonucleotide. For each molecule, three orthogonal views are provided. **D)** Plot of representative normalised Autocorrelation function vs correlation time observed in FCS experiment for 20nM Alexa-488 labeled CpGB (grey) and 20nM Alexa-488 labeled CpGB doped with 5µM of CpGB (red). They did not show any significant change in the diffusion time. Inset : The schematic diagram showing diffusion of fluorescent labeled oligonucleotide inside the confocal volume in FCS setup. **E)** Plot of representative normalised Autocorrelation function vs correlation time observed in FCS experiment for 20nM Alexa-488 labeled CpGA (grey) and 20nM Alexa-488 labeled CpGA doped with 5µM of CpGB (red). The arrow indicates a shift in the correlation function indicates self association. The arrow indicates the shift in the autocorrelation function. Inset : The schematic diagram showing diffusion of fluorescent labeled oligonucleotide inside the confocal volume in FCS setup**. F)** Plot of Number of particle (inside the confocal volume) vs. concentration of unlabeled oligonucleotide (CpGA in red and CpGB in black) observed in FCS experiment. CpGB did not show any significant change in the number of particle indicated range whereas CpGA showed a significant decrease in the number of particle which indicates self association. Three independent measurements were taken (n=3). The data were plotted as mean ± SD.

**Supplementary Figure 2: A)** Bar graphs demonstrating the uptake of labelled CpGA or CpGB by primary human pDCs treated with the indicated concentrations of pharmacological inhibitors targeting clathrin- and caveolin-mediated endocytosis- Pitstop 2 and Genistein, respectively (n= 7). Data is representative of at least three independent experiments. Student's t- test was used to calculate significance and data is represented as mean ± S.E.M. **B)** Dot plots showing knockdown efficiency of *ARHGAP26* mRNA expression in primary human pDCs, determined by qPCR (n=5). Data is represented as mean ± s.e.m and is representative of at least 3 independent experiments. Statistical significance was derived from student's t-test. **C)** Representative zebra plots demonstrating the uptake of CpGA-FITC or CpGB-FITC in control and ARHGAP26 KD primary human pDCs. **D)** Evolution of tension distribution in the population of cells after two minutes of treatment with CpGB/A compared with untreated condition. $n_{Cell\ (CpGA)}$=8, $n_{Cell\ (CpGB)}$=7 in each field; $n_{FBR\ (CpGA)}$ ~450, $n_{FBR(CpGB)}$ ~700. The Arrow indicates the population of regions from different cells that had increased tension on CPGA addition unlike CPGB addition.**E)** Evolution of $SD_{time}$ distribution in the population of cells after two minutes of treatment with CPGB/A compared with untreated condition. $n_{Cell\ (CpGA)}$=8, $n_{Cell\ (CpGB)}$=7 in each field; $n_{FBR\ (CpGA)}$ ~450, $n_{FBR(CpGB)}$ ~700.

**Supplementary Figure 3: A)** Schematic illustration of sequential correlative Total internal reflection Fluorescence - interference reflection microscopy imaging setup. **B)** Schematic representation of the work-flow followed to elucidate the local effect of oligonucleotide binding to plasma membrane of pDC cells. **C)** Representative whole field $SD_{time}$ maps of cells along time points for both CpGA treatment. Scale bar- 5 µm. For CpGA, $N_{cell}$: 9; $N_{cluster}$ : 46 . **D)** The radial profile of Fluorescence, Tension and $SD_{time}$ of a particular cell after 15 min of CpGA cargo addition that depicts a representative trend. **E)** Radial profiles of Fluorescence and

$SD_{time}$ around the Fluorescence peaks of CpGA clusters. Error denoted by the shaded region as SEM. * - represents significant change from the baseline condition for that particular time point after CpGA addition within 100 nm from centre. **F)** The spline-fitted probability distribution of the maximum surge in tension values from the cluster centre for all conditions from baseline to 5,15,20 min after CpGA addition. **G)** Radial profiles of Tension around the Fluorescence peaks of CpGA clusters for untreated condition and after 15 min of CpGA addition. Error denoted by the shaded region as SEM . * - represents significant change from the centre cluster tension for that particular time point through a distance of 325 nm from centre. Mann–Whitney U statistical significance test where ns- $p > 0.05$, * - $p < 0.05$, ** - $p < 0.001$. **H)** Dot plots showing knockdown efficiency of *PIEZO1* mRNA expression in primary human pDCs, determined by qPCR (n=11). Data is represented as mean ± s.e.m and is representative of at least 3 independent experiments. Statistical significance was derived from student's t-test.

**Supplementary Figure 4: A)** Schematic illustration of analysis followed for detection of fraction of actin on endosomal compartments. **B)** The illustration to corroborate that the cortical actin was disregarded and only internal actin patches were analyzed for estimating the actin distribution with endosomal compartments.

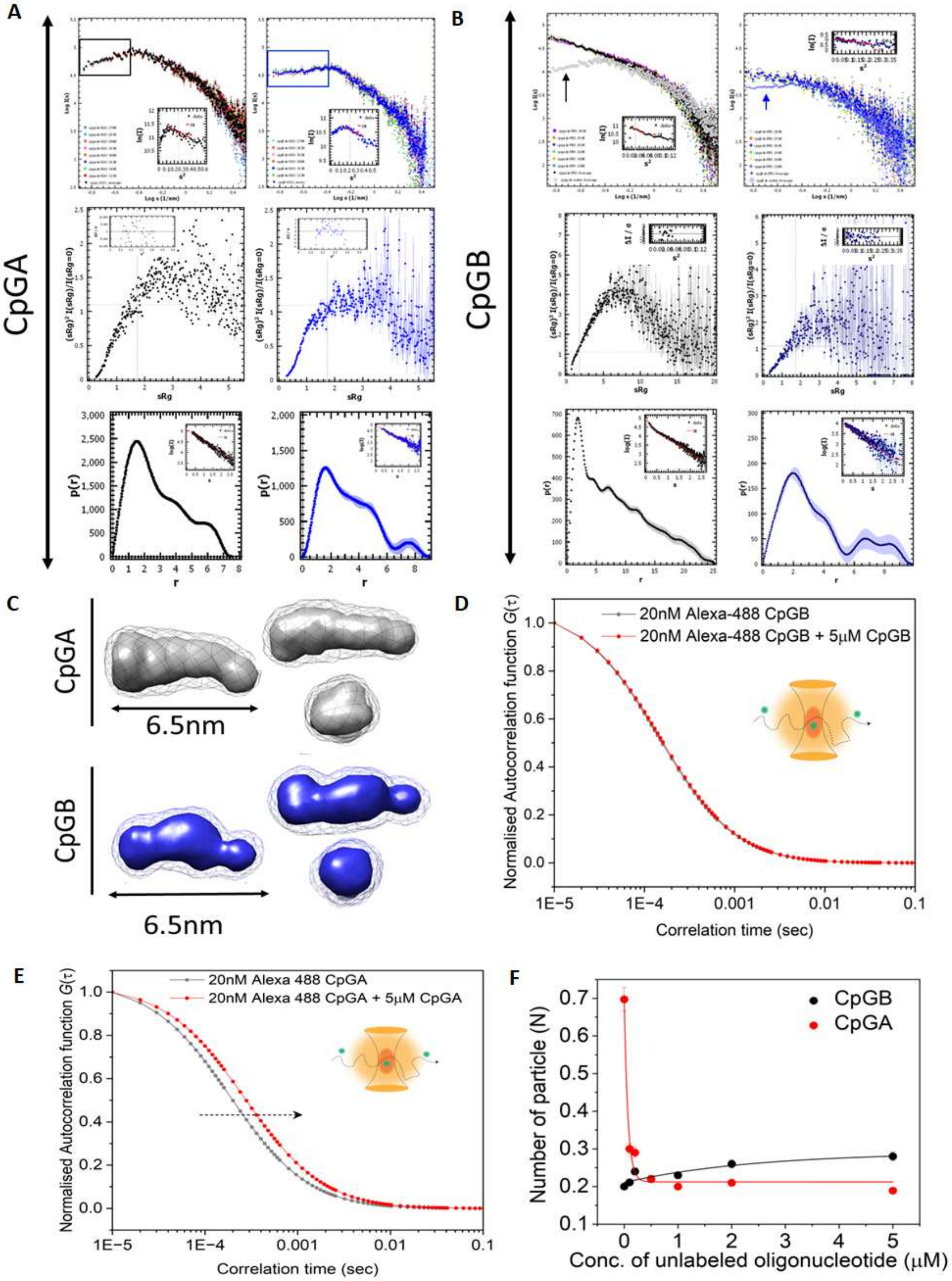

**Supplementary Figure 1**

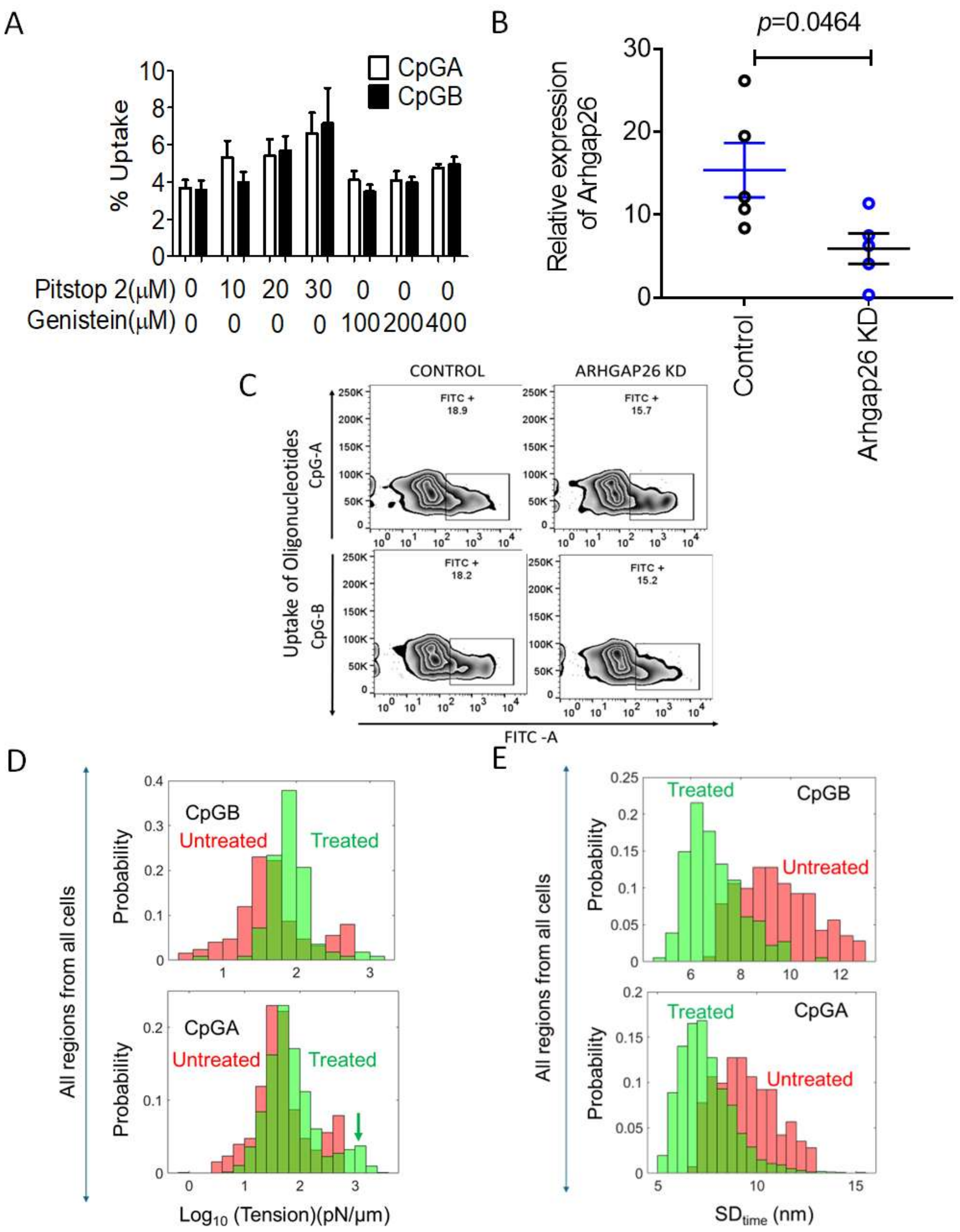

**Supplementary figure 2**

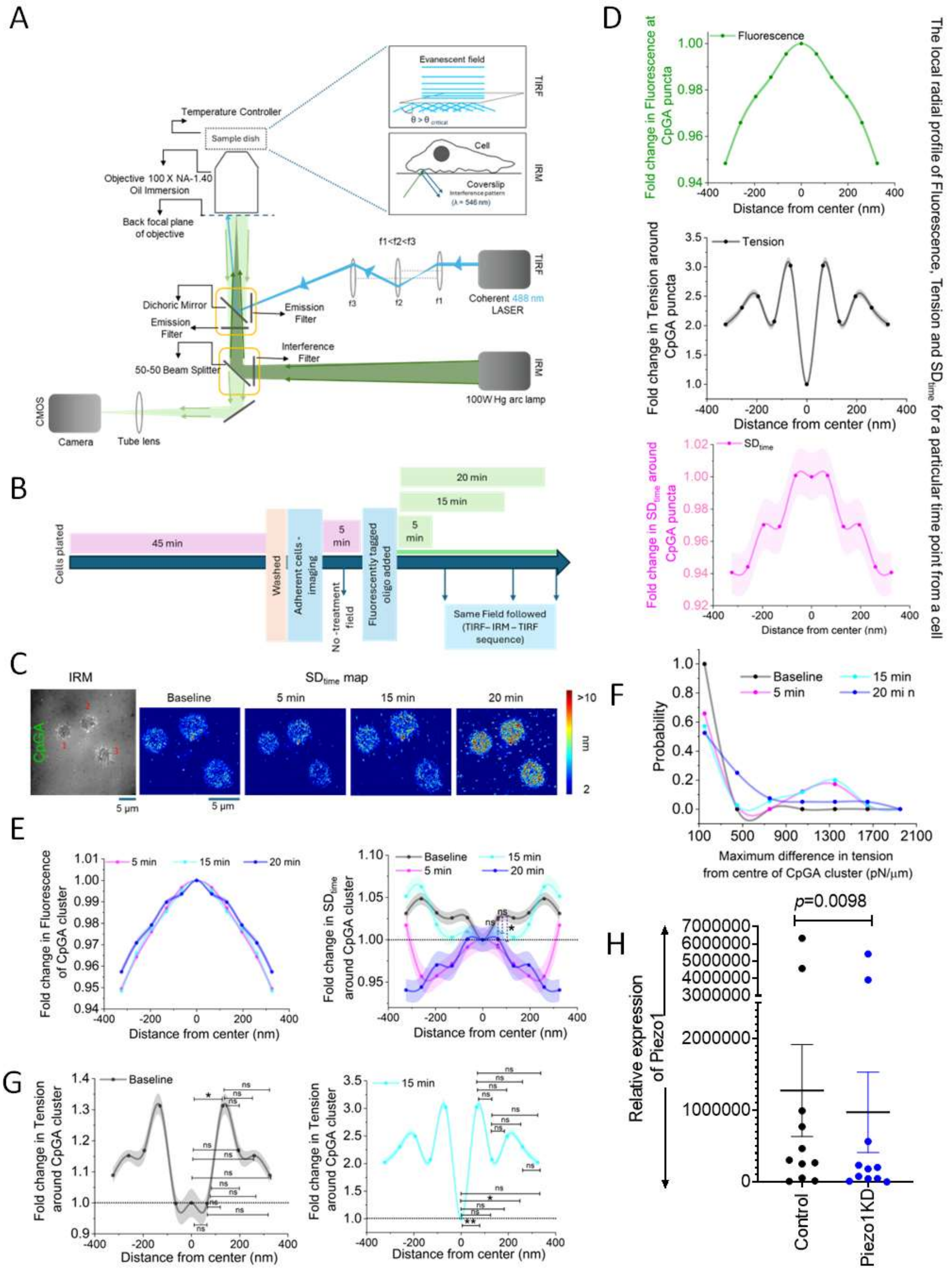

### Supplementary figure 3

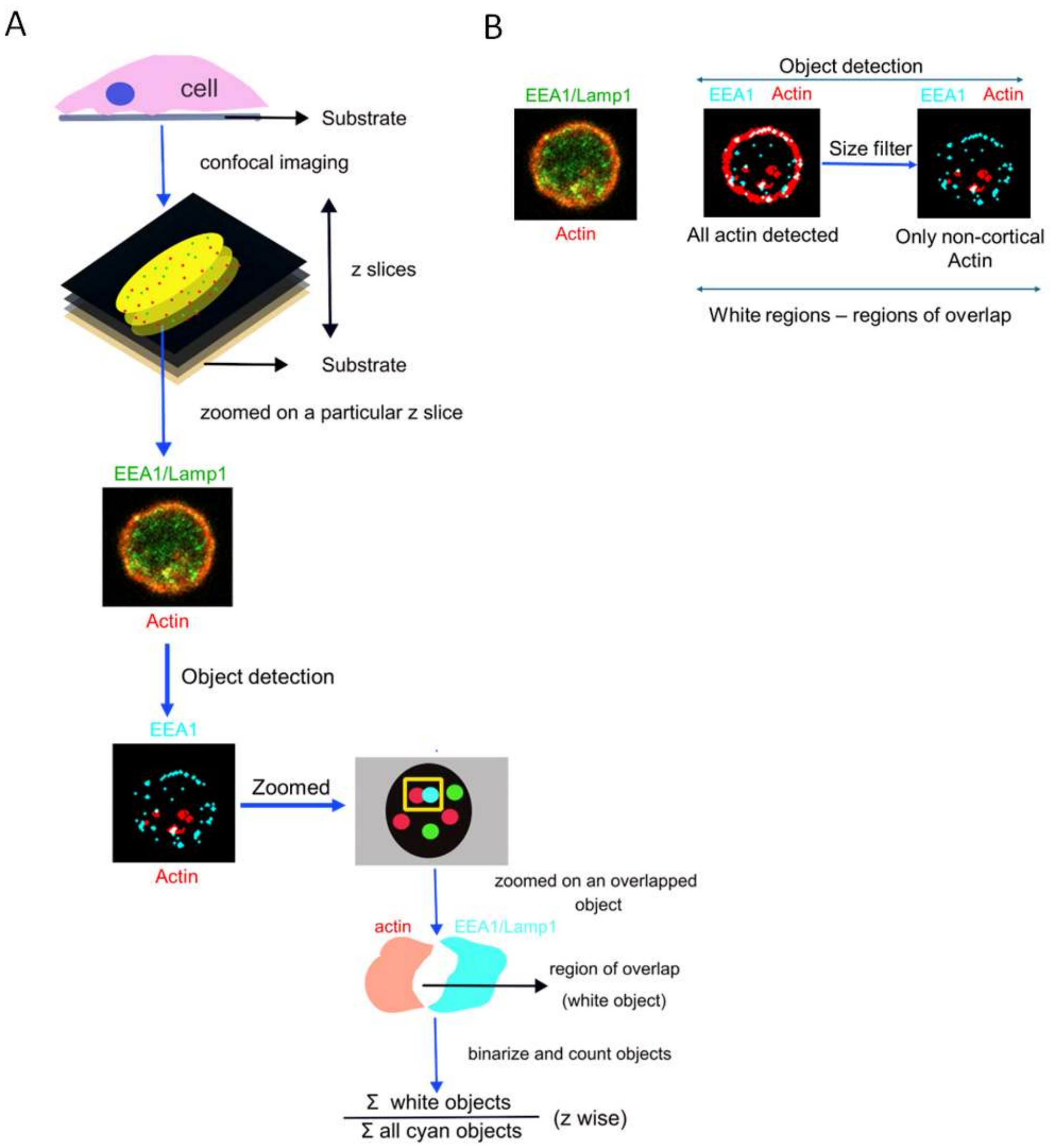

**Supplementary figure 4**

### Supplementary Videos 1-9

**Supplementary Video 1:** A typical field of fast timelapse IRM movie captured at 50 fps of adhered CpGA treated pDC, followed in time after 2 min CpGA addition, where each pixel is 72 nm. The time stamp indicates the duration of the movie. The scale bar represents 5 μm.

**Supplementary Video 2:** Representative 3D projection of Z- stacks of primary human pDCs stimulated with CpGA-FITC and stained with anti-EEA1 antibody and DAPI. The scale bar represents 3μm.

**Supplementary Video 3:** Representative 3D projection of Z- stacks of primary human pDCs stimulated with CpGA-FITC and stained with anti-LAMP1 antibody and DAPI. The scale bar represents 3 μm.

**Supplementary Video 4:** Representative 3D projection of Z- stacks of primary human pDCs treated with the Piezo1 inhibitor GsMTx4 for 30 minutess, followed by stimulation with CpGA-FITC and immunostaining with anti-EEA1 antibody and DAPI. The scale bar represents 3 μm.

**Supplementary Video 5:** Representative 3D projection of Z- stacks of primary human pDCs treated with the Piezo1 inhibitor GsMTx4 for 30 minutess, followed by stimulation with CpGA-FITC and immunostaining with anti-LAMP1 antibody and DAPI. The scale bar represents 3 μm.

**Supplementary Video 6:** Representative 3D projection of Z- stacks of primary human pDCs stimulated with CpGA, immunostained with anti-IRF7 and NFκB antibodies, and Hoechst for nuclear staining. The scale bar represents 3 μm

**Supplementary Video 7:** Representative 3D projection of Z-stacks of primary human pDCs treated with GsMTx4 for 30 minutes, followed by stimulation with CpGA and immunostaining with anti-IRF7 and anti-NFκB antibodies, as well as Hoechst for nuclear staining. The scale bar represents 3 μm.
